# Placental Nutrient Transport and Signaling in a Guinea Pig Model of Fetal Growth Restriction with Repeated Placental Nanoparticle-mediated IGF1 Treatment

**DOI:** 10.1101/2025.01.13.632768

**Authors:** Baylea N Davenport, Rebecca L Wilson, Alyssa A Williams, Helen N Jones

## Abstract

Roughly 10% of all pregnancies are affected by fetal growth restriction (FGR). The primary etiology of FGR is placental insufficiency: the placenta not providing the appropriate amount of nutrients and oxygen to the fetus. There is currently no treatment for FGR or placental insufficiency. Because of the placentas pivotal role in FGR and supplying nutrients to the fetus, it offers an excellent target for therapeutic intervention. Using a guinea pig maternal nutrient restriction model and a repeated placental nanoparticle-mediated IGF1 treatment, placental IGF1 signaling and nutrient transport pathways were characterized to understand changes with FGR and treatment. This study elucidates the signaling mechanisms in which repeated placental nanoparticle-mediated IGF1 treatment leads to correct fetal growth. Overall, this study resulted in sex-specific kinase signaling and nutrient transporter changes within the placenta in both FGR and treatment groups. Combined with our previous studies using this treatment, we demonstrate the basic molecular signaling of this treatment and recapitulate the plausibility of this therapy for future human translation.

## INTRODUCTION

Fetal growth restriction (FGR) is characterized by a fetus failing to reach their intrauterine growth potential [1, 2]. While perinatal outcomes are largely influenced by severity, FGR is the leading cause of preterm delivery and stillbirth [3]. Adequate growth *in utero* is critical for fetal development and is a crucial contributor to long-term health. When not victim to stillbirth, those affected by FGR are more likely to develop health and developmental complications such as metabolic, cardiovascular or neurodevelopmental disorders [4]. One of the most common contributors to FGR is the placenta failing to transfer adequate nutrients and oxygen to the fetus, known as placental insufficiency [5, 6].

Fetal growth is primarily controlled through maternal circulating nutrients such as glucose and amino acids transported across the placenta into fetal circulation [7–9]. Inadequate supply of nutrients to the fetus can be multi-factorial. While availability of nutrients in the maternal circulation is imperative, transport from maternal to fetal circulation is not passive, thus if the placenta itself is insufficient, this can also lead to further decreased nutrients in fetal circulation [9, 10]. Placental insufficiency can be multifold with both functional and structural deficits. Syncytiotrophoblast membranes express glucose and amino acid transporters for active facilitation from maternal to fetal circulation [9]. Human cases of FGR have identified varying activity and expression levels of these transporters with deficiency compared to control placentas [8, 9, 11]. Changes in placenta structure with FGR have also been found to decrease nutrient and oxygen delivery to the fetus. Deficiencies in utero-placental blood flow, vascular exchange area size, and interhaemal distance have all been associated with decreased nutrient transfer and FGR [10, 12–14]. The placenta’s active role in nutrient transport is one of the many reasons we choose to target this organ for intervention in FGR.

While there are currently no treatments for FGR in the human, there is a growing effort for potential therapies [7, 15–23]. We have developed a nanoparticle-mediated, placenta targeted gene therapy to deliver IGF1 for the treatment of placental insufficiency and FGR [15]. IGF1 is a master regulator with key roles in angiogenesis and nutrient transport in the placenta throughout the entirety of gestation and is known to be downregulated in FGR placentas [24–26]. To model FGR we employ the guinea pig maternal nutrient restriction model: a well characterized animal model of placental and fetal development that establishes placental insufficiency and FGR [8, 27–31]. Complexing a non-viral copolymer with an *hIGF1* plasmid under a trophoblast specific promoter, we initially performed a single ultrasound guided intraplacental injection at mid-pregnancy after FGR has been established. 5 days post-injection we found an increase in glucose and amino acid transporters within the placenta and an increase in circulating fetal glucose [32]. These short-term improvements to nutrient transport led to us now performing repeated treatment (3 times over the latter half of pregnancy) where we have shown corrected fetal glucose levels, improved placental structure, and corrected fetal growth [33].

While we have shown corrected fetal growth with repeated IGF1 treatment, here we aim to show the underlying mechanism of increased placental *hIGF1* leading to this improved outcome, as well as elucidating any sexually dimorphic signaling mechanisms. Human studies of FGR have repeatedly shown sexually dimorphic responses to in-utero insults and post-natal NICU outcomes, with male fetuses having poorer outcomes than their female counterparts [34, 35]. Our studies and others in the field have previously reported similar outcomes in animal models and aim to understand the molecular and physiological mechanisms leading to these sexual dimorphisms in FGR and with IGF1 treatment. To further understand the mechanisms of nanoparticle mediated *hIGF1* treatment that leads to improved placental function and fetal growth, we sought to identify the pathways influenced by this therapy in the FGR placenta that lead from IGF1 delivery to increased fetal growth through kinase signaling and nutrient transporter function.

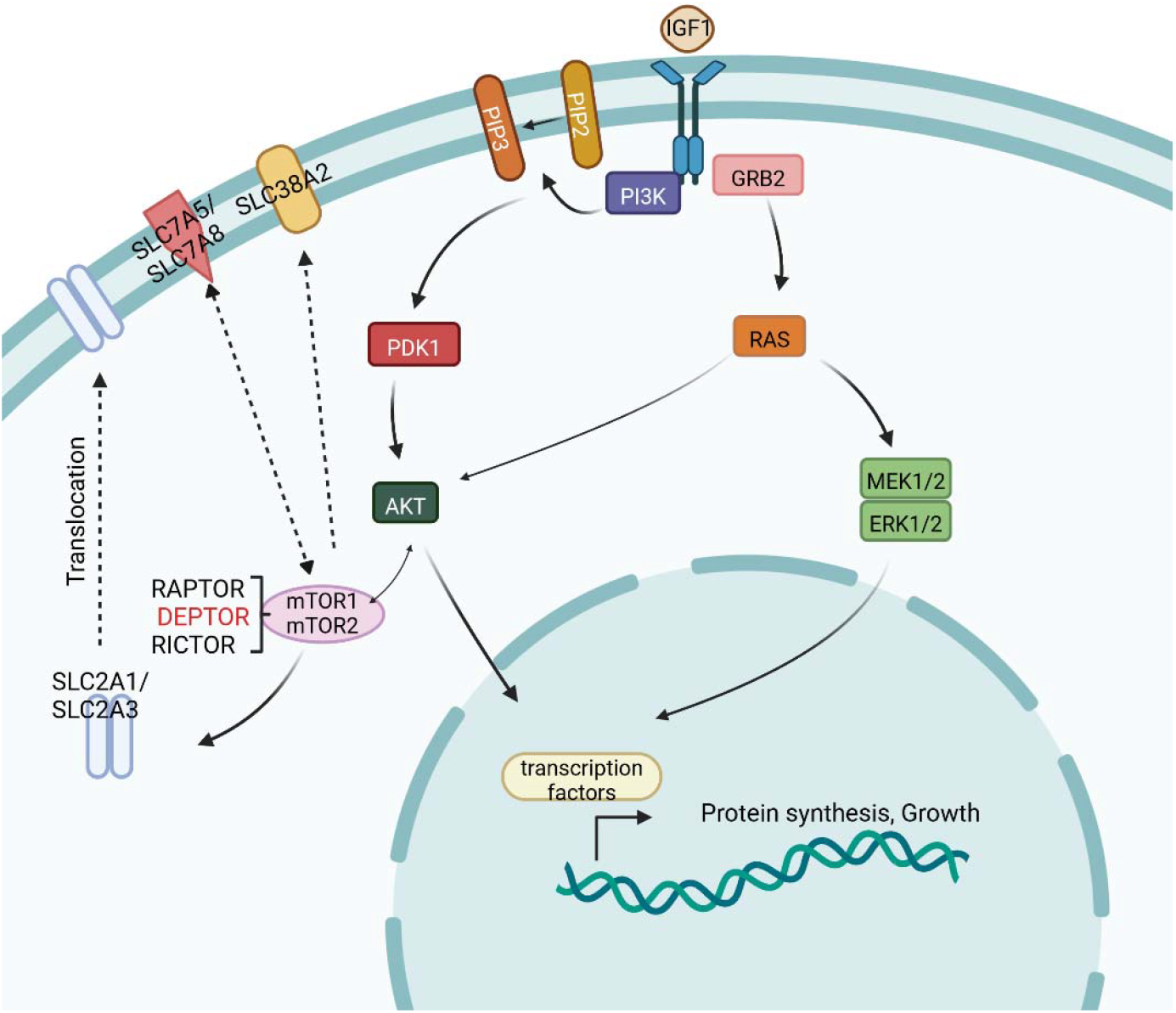

## METHODS

### Nanoparticle Synthesis

Nanoparticles were formed by complexing a lyophilized non-viral PHPMA_115_-b-PDMEAMA_115_ co-polymer reconstituted in water with plasmid containing the *hIGF1* gene under the control of a trophoblast-specific promoter, *CYP19A1*. 50 μg of plasmid was suspended in a 200 μL volume solution at room temperature. Detailed methods of copolymer synthesis and nanoparticle formation can be found in Wilson et al., 2022 [32].

### Animal Husbandry, Mating, and Intraplacental Nanoparticle Injections

Animal care and usage was approved by the Institutional Animal Care and Usage Committee at the University of Florida (Protocol #202011236). Female Dunkin-Hartley guinea pigs (Dams) were purchased from Charles River Laboratories (Wilmington, MA) at 500–550 g (~ 8-9 weeks of age). Animals were housed in an environmentally controlled room (22°C/72°F, 50% humidity 12 h light-dark cycle). Upon arrival, food (LabDiet diet 5025: 27% protein, 13.5% fat, and 60% carbohydrate as % of energy) and water were provided ad libitum. Guinea pigs were acclimatized for 2 weeks prior to being assigned to either ad libitum diet (termed Control: n = 6) or maternal nutrient restriction (MNR) diet (n=12). Assignment was done by ranking animals heaviest to lightest and systematically assigning them to each group for even biological weight distribution. MNR diet consists of a 70% food intake diet based on kilogram of body weight of control from 4 weeks pre-mating through mid-pregnancy (GD35), then increased to 90% food intake through term to maintain pregnancy. Precise details on timed matings, pregnancy confirmation, and ultrasound-guided nanoparticle injections can be found in Wilson, et al 2022 and Davenport, et al 2024 [32, 33]. Nanoparticle delivery was performed a total of 3 times, each 8 days apart from mid pregnancy till term (GD36,44,52±3) with guinea pigs being sacrificed 8 days after the final injection (GD60±3) near term (guinea pig gestation is 65-70 days). One placenta per litter was injected with either nanoparticle mediated *hIGF1* (MNR+IGF1 n=6) or a non-expressing sham nanoparticle (Control n=6, MNR n=6). In the MNR+IGF1 dams, placentas were separated based on receiving a direct injection of nanoparticle-mediated *hIGF1* “MNR+IGF1 (Direct Injection)” or by being indirectly exposed to circulating residual nanoparticle-mediated *hIGF1* “MNR+IGF1 (Indirect Exposure)” which was further confirmed by expression levels of *hIGF1* in each placenta via qPCR [33]. Dams were sacrificed via carbon dioxide asphyxiation followed by cardiac puncture and exsanguination. Fetuses (Control: n=8 female and n=11 male, MNR: n=5 female and 11 male, MNR + IGF1: n=6 female and 10 male) and the delivered placenta (placenta, sub-placenta, and decidua) were removed from the uterus and weighed. Fetal sex was determined at this time by examination of the gonads. Here it was determined that by random chance only male fetuses received direct injections (as we cannot determine fetal sex at time of injection). Placentas were processed for histology (fixed in 4% PFA) or molecular analysis (preserved in RNAlater and snap-frozen). To reduce bias, all subsequent analyses were performed blinded. For in-depth synopsis of all mating, pregnancy, litter size, statistics, etc see Davenport, Wilson, et al 2024 [33].

### Quantitative PCR

Approximately 150–200 mg of placenta labyrinth tissue preserved in RNAlater was lysed and homogenized in RLT-lysis buffer (Qiagen) using a tissue homogenizer. RNA was extracted and DNase treated using the RNeasy Midi Plus Kit (Qiagen) following standard manufacturers protocols. 1 µg of RNA was used to convert to cDNA using the High-capacity cDNA Reverse Transcription kit (Applied Biosystems), following standard manufacturers protocol. qPCR was performed in a 20 μL reaction using 2.5 μL of cDNA, 10 μL of PowerUP SYBR green (Applied Biosystems), 1.2 μL of 10 nM forward and reverse primers, and 5.1 μL of water. Primer sequences of all genes can be found in Supplemental table 2. All gene expression was normalized to the mean of reference genes *B-actin* and *Rsp20*. Reactions were performed using the Quant3 Real-Time PCR system (Applied Biosystems). Relative mRNA expression was calculated using the comparative CT method with the Design and Analysis Software v2.6.0 (Applied Biosystems).

### Western Blots

Snap frozen placenta tissue was homogenized in ice-cold RIPA buffer with protease and phosphatase inhibitors (Thermo Fisher). A Bradford protein assay was performed to determine protein concentrations (Pierce Coomassie Plus, Thermo Fisher) following manufacturer’s protocol. 40μg of protein was combined with SDS loading buffer and reducing agent (Invitrogen) and then denatured at 95°C for 10 minutes. Protein lysates were loaded into 4-12% tris-bis bolt pre-cast gels (Invitrogen) and run for 1 hour at 150 volts. Protein was transferred to nitrocellulose membranes at 30 volts for 1.5 hours on ice. Protein transfer was confirmed with Ponceau S staining before blocking in 5% skim-milk in TBST (Tris-buffered saline containing Tween20) overnight at 4°C. Primary antibodies (Supplemental table 1) were diluted in 5% skim-milk in TBST and applied to membranes for 2 hours at room temperature. Membranes were washed for 5 minutes, 3 times in TBST, then placed in HRP-conjugated secondary antibody (Cell Signaling 7074 or 7076, 1:1000) for 2 hours. Membranes were washed once again before visualizing bands using SuperSignal West Femto Maximum Sensitivity Substrate (Thermo Fisher) on a Chemidoc Imager (*Bio*-*Rad*). Signal intensity of the protein bands was calculated using Image Lab software (version 6.1, *Bio-Rad*) and normalized to β-actin.

### Immunohistochemistry

PFA fixed placenta tissue was processed and paraffin embedded. 5 μm thick sections were cut and slide-mounted for immunohistochemistry (IHC). Placenta sections were de-waxed and rehydrated using Histo-clear and ethanol following standard protocols. Antigen retrieval was then performed by incubating slides in Target Retrieval Solution (Invitrogen) for 20 minutes at 95°C, then cooled to room temperature for an additional 20 minutes. Endogenous peroxidases were blocked using 3% hydrogen peroxide for 10 minutes before blocking in animal free protein block (Vector) for 30 minutes at room temperature. Primary antibodies (Supplemental Table 1) were diluted in protein block and incubated overnight at 4°C. Slides were then washed in PBS for 5 minutes, 3 times before adding a biotinylated secondary antibody (Invitrogen, BA-9200 or BA-1000) at 1:200 dilution. Slides were once again washed before incubating in ABC (avidin-biotin complex) reagents (Vector) and visualized with DAB staining (3,3′-diaminobenzidine) (Vector). Nuclei were counterstaining using hematoxylin and coverslips mounted using DPX mounting solution (Millipore). Sections were imaged using the Zeiss Axioscan Scanning Microscope at 40x magnification. Images were obtained through the Zen Imaging Software where 10 region of interest images were taken of 75×100μm labyrinth sections. Intensity and localization were qualitatively measured by 2-3 blinded lab members on staining intensity with localization being in the nucleus, cytoplasm, membrane, or non-specific.

### Statistical Analysis

Statistical analyses were performed using SPSS Statistics v.29. Distribution assumptions were checked with a Q-Q-Plot. Statistical significance was determined using generalized estimating equations with gamma as the distribution and log as the link function. Dams were considered the subject, diet and nanoparticle-mediated *IGF1* treatment treated as main effects, and gestational age and litter size as covariates. In the sham treated Control and MNR groups, there was no effect of direct placental injection for any outcomes measured and was therefore removed as a main effect in these groups. Placentas from female and male placentas were analyzed separately. In male fetuses, direct injection or indirect exposure of nanoparticle-mediated *IGF1* was also considered as a main effect. Statistical significance was considered at P≤0.05. For statistically significant results, a Bonferroni post hoc analysis was performed. Results are reported as estimated marginal means ± 95% confidence interval.

## RESULTS

Results on maternal and fetal weight/physiological outcomes as well as placental weight and efficiency have been previously published [33]. No adverse health outcomes for mother or fetus were found. At time of sacrifice it was determined that all placentas that received repeated direct injection of *hIGF1* nanoparticle treatment were male. Because we cannot determine sex at time of injections and this random determination occurred, all reported data for males have both MNR+IGF1 (Direct Injection) and MNR+IGF1 (Indirect Exposure) (supplemental data) where females only include MNR+IGF1 (Indirect Exposure). Repeated *hIGF1* nanoparticle treatment resulted in placental expression of *hIGF1*, with placentas that received direct injection displaying higher levels of *hIGF1* than littermates that were indirectly exposed [33]. Near-term fetal weight was restored from MNR with *hIGF1* nanoparticle treatment back to Control levels in both sexes, as well as aberrant glucose levels in MNR also corrected with *hIGF1* nanoparticle treatment back to Control levels [33]. Delivered placental weight and efficiency did not change in males, while placental efficiency decreased with MNR females but was restored back to Control levels with *hIGF1* nanoparticle treatment [33].

In placentas of both male and female placentas, protein expression of total ERK (Extracellular signal-regulated kinase) did not change between any groups at end of term (Figure 1A, B, Supp.1A). In males, protein expression of phosphorylated ERK (pERK, p44,42, Thr202/Tyr204) (phosphorylated/total ERK expression) did not significantly change among groups but trended downward in MNR+IGF1 (Direct Injection) placentas compared to MNR (p=0.057) (Figure 1C, Supp.1B). In females, pERK expression did not change between any groups (Figure 1D). In males, while not statistically significant, total AKT (also known as Protein kinase B) expression decreased near significance in MNR compared to Control (p=0.067) (Figure 1E, Supp. 1C). AKT expression also trended upward in MNR+IGF1 (Direct Injection) placentas compared to MNR with near significance (p=0.053), with expression levels similar to Control in males. In placentas of female fetuses, protein expression of total AKT did not change between groups (Figure 1F). Phosphorylated AKT (pAKT, Ser473) expression in males significantly decreased in MNR+IGF1 (Direct Injection) placentas compared to MNR (p<0.05) (Figure 1G, Supp. 1D). pAKT expression had near significant increase in MNR compared to Control placentas in females but not statistically significant (p=0.06) (Figure 1H).

**Figure 1.**
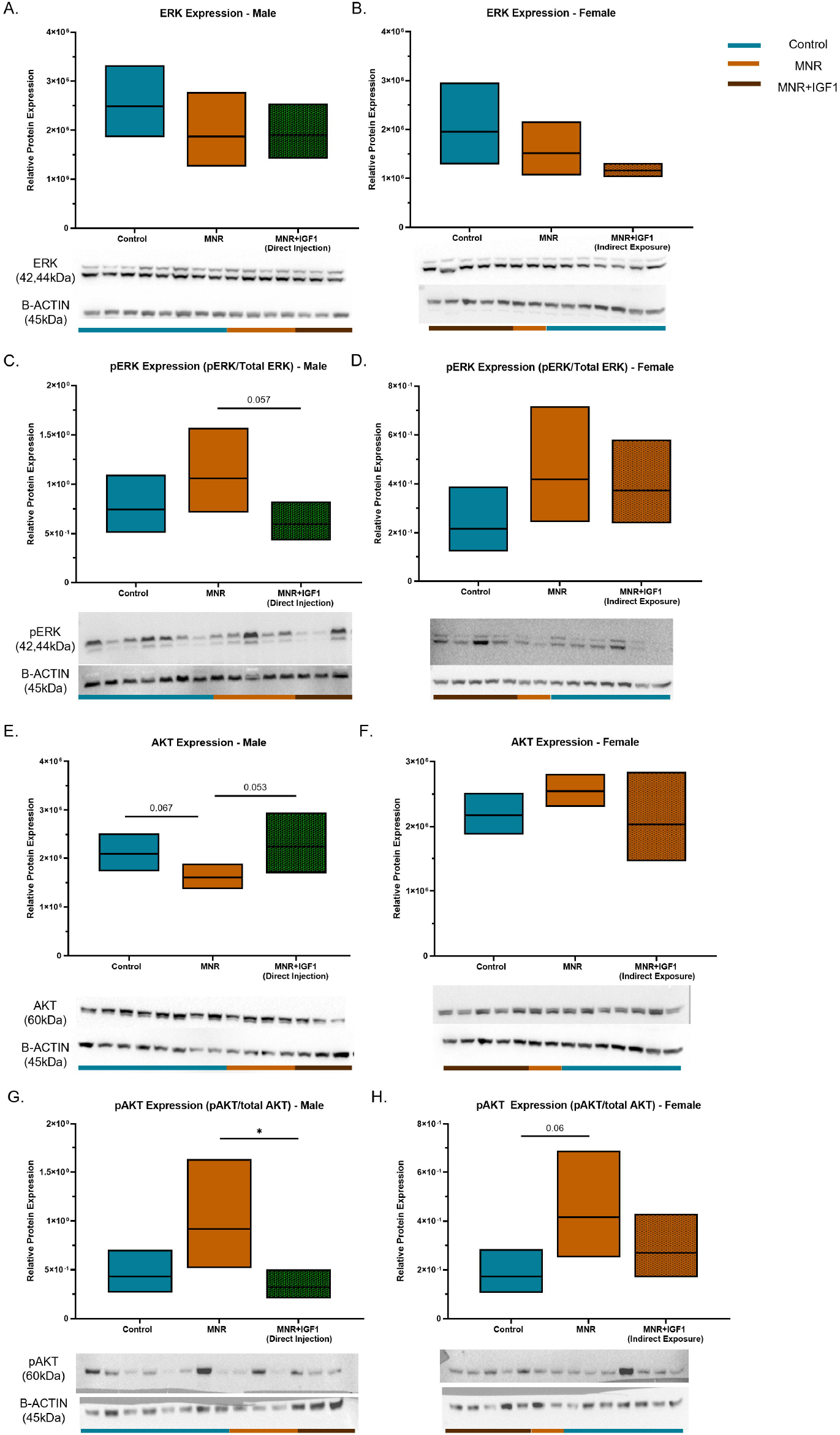
Effects of maternal nutrient restriction (MNR) and repeated *hIGF1* nanoparticle gene therapy (MNR+*IGF1*) on placental kinase signaling at end of term. **A, B**. In placentas of male and female fetuses, there was no difference in total ERK protein expression between all groups. **C**. pERK protein expression in MNR+IGF1 (Direct Injection) placentas had a near significance decrease from MNR placentas in males, but did not reach statistical significance. **D**. pERK protein expression did not change between groups in placentas of female fetuses. **E**. Total AKT expression had a near significance decrease in MNR placentas compared to Control, while MNR+IGF1 (Direct Injection) placentas trended back up from MNR, comparable to Control in placentas of male fetuses. **F**. There were no differences in total AKT protein expression among females. **G**. pAKT protein expression decreased in the MNR+IGF1 (Direct Injection) placentas compared to MNR in males. **H**. In females, pAKT expression had a near significant increase in MNR placentas compared to Control, but did not reach statistical significance. Control (+ sham treatment): n = 6 dams (8 female and 11 male fetuses), MNR (+ sham treatment): n = 6 dams (5 female and 11 male fetuses), MNR + IGF1: n = 5 dams (6 female and 10 male fetuses). Western blot images show representative blots and do not include all blots/samples included in analysis. Samples were normalized across blots. Colored bars below western blots indicate sample group: teal=Control, orange= MNR, maroon= MNR+IGF1. Data are estimated marginal means ± 95% confidence interval. P values calculated using generalized estimating equations with Bonferroni post hoc analysis. *P≤0.05; **P≤0.01, ***P≤0.001

There were no changes in protein expression of total mTOR (mammalian target of rapamycin) in male or female placentas (Figure 2A, B). Phosphorylated mTOR (p-mTOR) increased in MNR+IGF1 (Indirect exposure) compared to Control and MNR (p<0.001, p<0.001) in male placentas (Figure 2C, Supp. 1F). Conversely, in females, p-mTOR decreased in MNR and MNR+IGF1 (Indirect Exposure) placentas compared to Control (p<0.05) (Figure 2D). There were no changes in RICTOR (RPTOR Independent Companion of MTOR Complex 2) protein expression among any groups of either sex (Figure 2E, F, Supp. 1G). In males there were no changes in RAPTOR (Regulatory-associated protein of mTOR) expression between groups (Figure 2G, Supp. 1H). In females, RAPTOR expression decreased in MNR+IGF1 (Indirect Exposure) placentas compared to MNR (p<0.05) (Figure 2H). In males, DEPTOR (DEP domain-containing mTOR-interacting protein) expression in both MNR+IGF1 (Indirect exposure) (Control p<0.001, MNR p<.01) and MNR+IGF1 (Direct Injection) (Control p<0.01, MNR p<.01) placentas decreased compared to Control and MNR placentas (Figure 2I, Supp. 1I). Similarly, in females, DEPTOR expression decreased in MNR+IGF1 (Indirect Exposure) placentas compared to Control (p<0.05) and trended downward from MNR (p=0.056) (Figure 2J).

**Figure 2.**
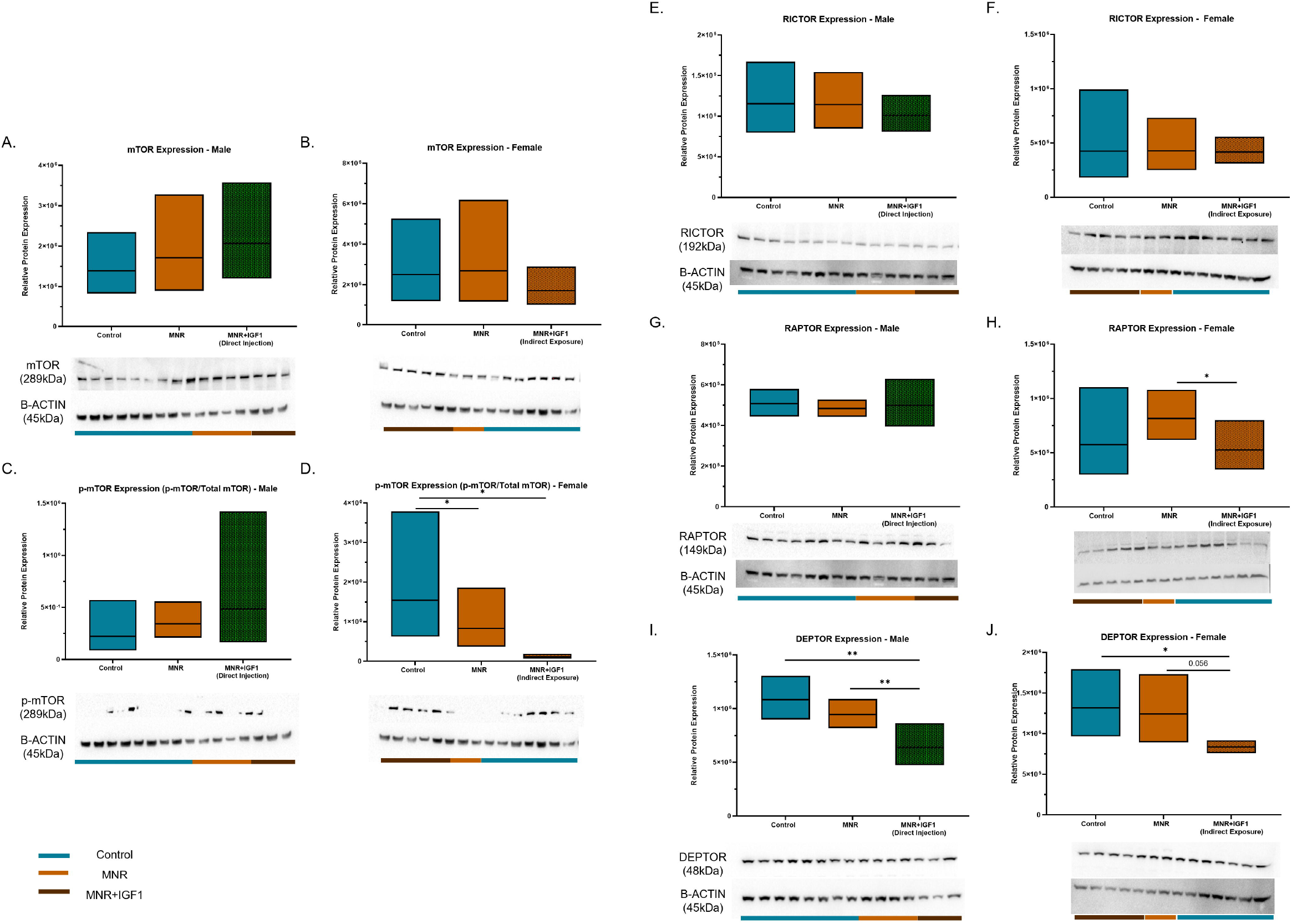
Effects of maternal nutrient restriction (MNR) and repeated *hIGF1* nanoparticle gene therapy (MNR+*IGF1*) on placental mTOR signaling at end of term. **A, B**. Total mTOR protein expression was unaltered among all groups in placentas of male and female fetuses. **C**. In males p-mTOR protein expression was unaltered between groups. **D**. In females, p-mTOR protein expression decreased in both MNR and MNR+IGF1 (Indirect exposure) placentas compared to Control. **E, F**. No groups among males or females expressed differences in placental RICTOR protein expression. **G**. RAPTOR expression in male placentas did not change across groups. **H**. RAPTOR protein expression decreased in MNR+IGF1 (Indirect Exposure) placentas compared to MNR in females. **I**. DEPTOR protein expression was significantly decreased in MNR+IGF1 (Direct Injection) compared to both Control and MNR in males. **J**. In females, DEPTOR protein expression was significantly decreased in MNR+IGF1 (Indirect Exposure) compared to Control. Control (+ sham treatment): n = 6 dams (8 female and 11 male fetuses), MNR (+ sham treatment): n = 6 dams (5 female and 11 male fetuses), MNR + IGF1: n = 5 dams (6 female and 10 male fetuses). Western blot images show representative blots and do not include all blots/samples included in analysis. Samples were normalized across blots. Colored bars below western blots indicate sample group: teal=Control, orange= MNR, maroon= MNR+IGF1. Data are estimated marginal means ± 95% confidence interval. P values calculated using generalized estimating equations with Bonferroni post hoc analysis. *P≤0.05; **P≤0.01, ***P≤0.001

There was no change in total 4EBP1 protein expression among any group of either sex (Figure 3A,B). Placentas of male fetuses also had no change in phosphorylated 4EBP1 (p-4EBP1) (Figure 3C). In females, however, p-4EBP1 decreased in MNR (p<0.001) and MNR+IGF1 (Indirect Exposure) (p<0.001) placentas compared to Control (Figure 3D).

**Figure 3.**
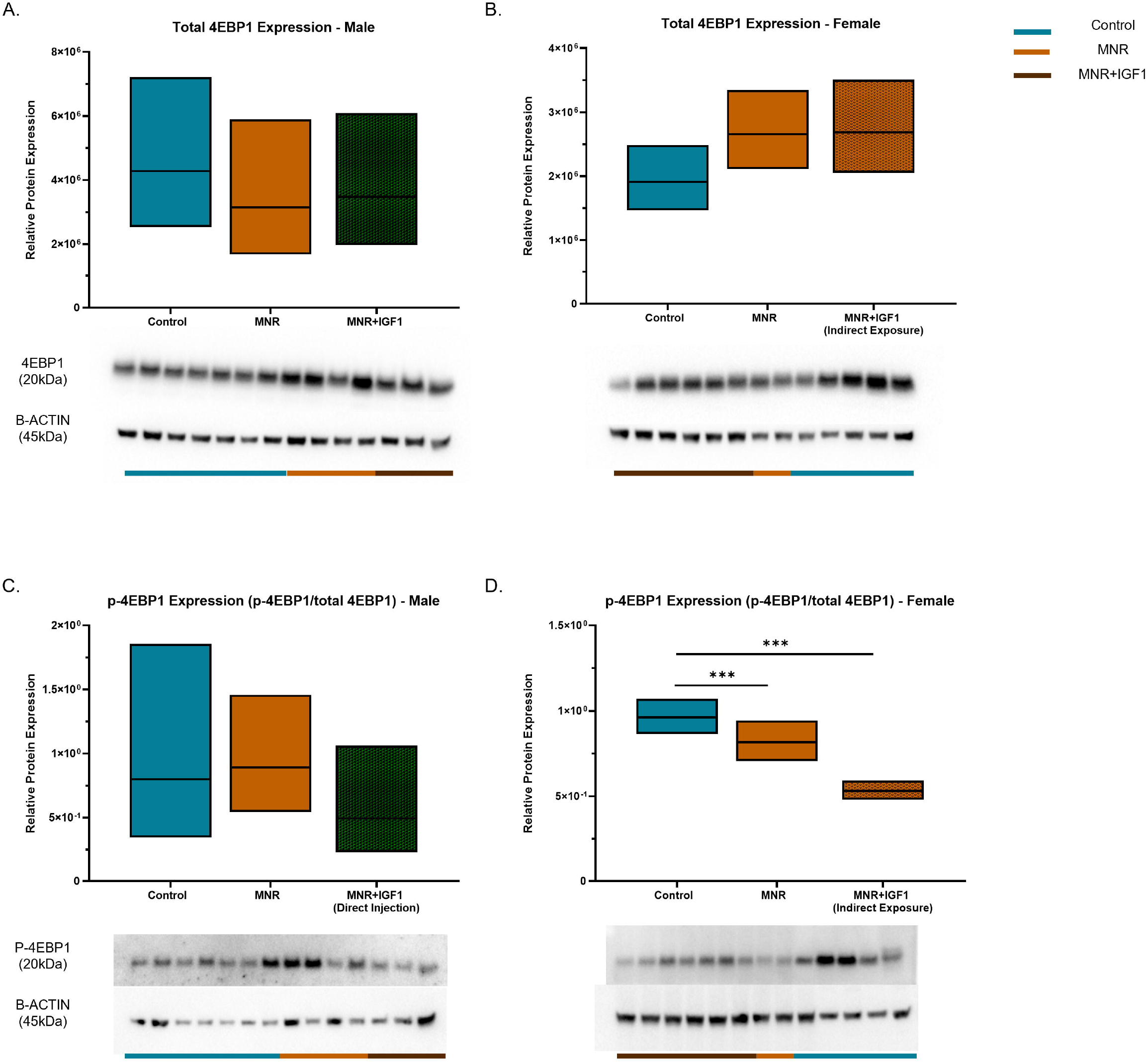
Effects of maternal nutrient restriction (MNR) and repeated *hIGF1* nanoparticle gene therapy (MNR+*IGF1*) on placental downstream mTOR signaling at end of term. **A, B**. Total 4EBP1 protein expression did not change among any groups in placentas of male or female fetuses. C. In males, p-4EBP1 protein expression did not change among groups. D. In females, p-4EBP1 protein expression decreased in both MNR and MNR+IGF1 (Indirect exposure) placentas compared to Control. Control (+ sham treatment): n = 6 dams (8 female and 11 male fetuses), MNR (+ sham treatment): n = 6 dams (5 female and 11 male fetuses), MNR + IGF1: n = 5 dams (6 female and 10 male fetuses). Western blot images show representative blots and do not include all blots/samples included in analysis. Samples were normalized across blots. Colored bars below western blots indicate sample group: teal=Control, orange= MNR, maroon= MNR+IGF1. Data are estimated marginal means ± 95% confidence interval. P values calculated using generalized estimating equations with Bonferroni post hoc analysis. *P≤0.05; **P≤0.01, ***P≤0.001

In males, *Glut1* (*Slc2a1*) RNA expression had a near significant increase in MNR+IGF1 (Indirect Exposure) placentas compared to Control (p=0.054) and was significantly increased compared to MNR (p<0.001) (Figure 4A, Supp. 2A). GLUT1 protein expression in males showed no significant changes by western blot, though trended down in MNR compared to Control and MNR+IGF1 (Direct Injection) (p=0.09, p=0.09) (Figure 4B, Supp. 2B). IHC demonstrates similar results with localization of GLUT1 showing defined membrane staining in Control and MNR+IGF1 (Direct Injection) placentas, but much more diffuse staining not localized to the membrane in MNR comparably (Figure 4C). In females, *Glut1* RNA expression decreased in MNR+IGF1 (Indirect Exposure) placentas compared to Control (p<0.05) (Figure 4D), however, protein expression fif not change between groups by western blot (Figure 4E). Localization, however, showed similar results to males with defined membrane staining in Control, less defined membrane staining in MNR, and MNR+IGF1 (Indirect Exposure) more similar to Control (Figure 4F). In males, *Glut3* (*Slc2a3*) RNA expression increased in MNR+IGF1 (Direct Injection) placentas compared to MNR (p<0.05) (Figure 4G, Supp. 2C). Protein expression of GLUT3 decreased in males in MNR+IGF1 (Indirect Exposure) compared to Control (p<0.05) (Figure 4H, Supp. 2D). In females, *Glut3* RNA nor GLUT3 protein expression/localization had significant changes (Figure 4J,K,L). There were also no changes in *Glut8* (*Slc2a8*) or *Glut9* (*Slc2a9*) expression among any groups of either sex (Supp. 3A-D).

**Figure 4.**
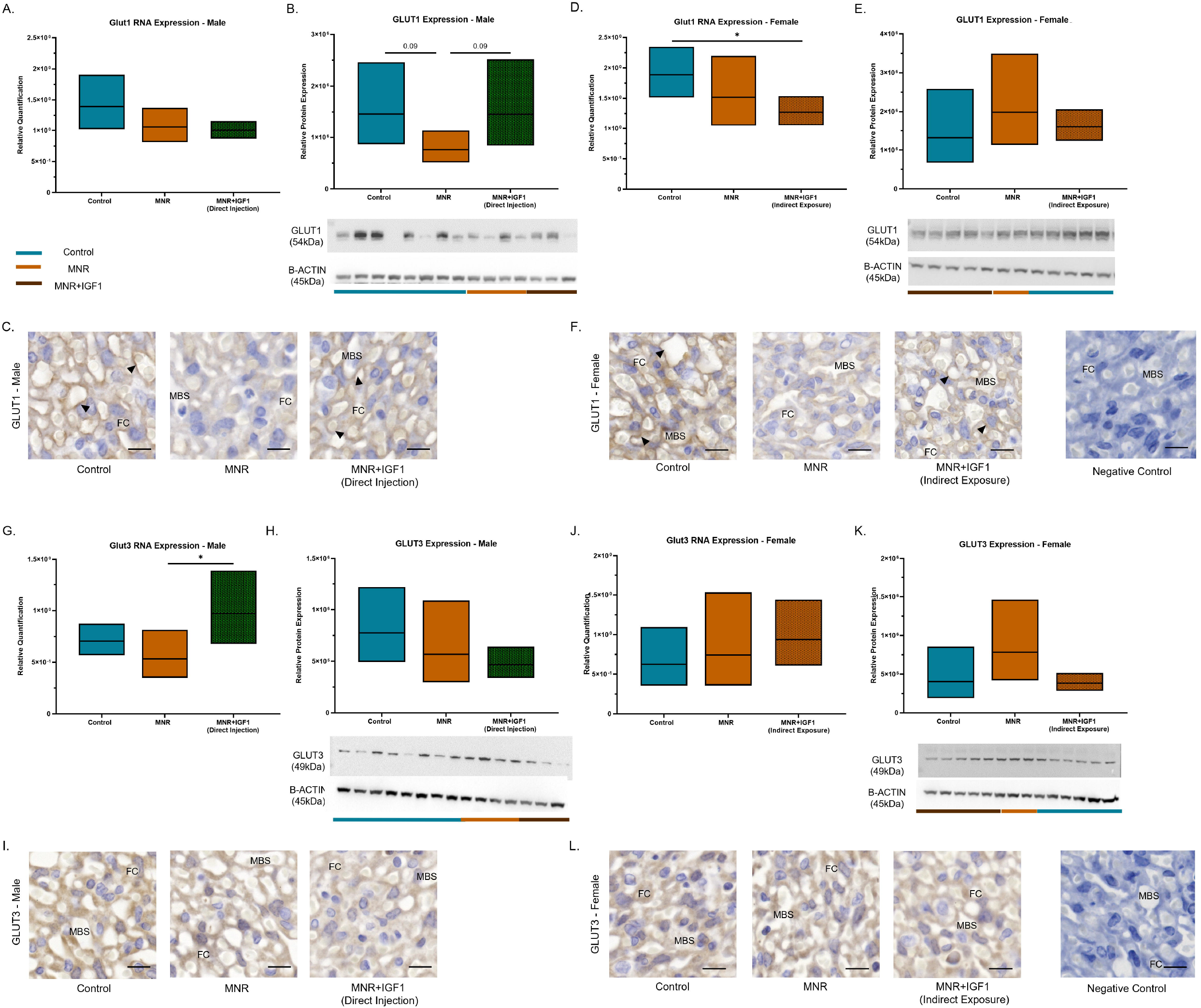
Effects of maternal nutrient restriction (MNR) and repeated *hIGF1* nanoparticle gene therapy (MNR+*IGF1*) at end of term on placental glucose transporters. **A**. In male placentas *Glut1* mRNA expression did not change among groups **B**. GLUT1 protein expression did not change among groups in male placentas. **C**. GLUT1 IHC showed defined, pronounced membrane staining in Control, more diffuse/lighter staining in MNR, and similar staining in MNR+IGF1 (Direct Injection) to Control in males. **D**. In females, *Glut1* placental mRNA expression decreased in MNR+IGF1 (Indirect Exposure) compared to controls. **E**. There was no change in GLUT1 protein expression in female placentas. **F**. In females, GLUT1 IHC showed defined, pronounced membrane staining in Control. MNR showed more diffuse/lighter staining, and a return in membrane staining in MNR+IGF1 (Indirect Exposure) compared to MNR, more similar to Control. **G**. Male *Glut3* mRNA expression increased in MNR+IGF1 (Direct Injection) compared to MNR placentas. **H**. GLUT3 protein expression did not change between groups in males. **I**. Localization of GLUT3 showed no changes among males. **J**,**K**,**L**. In females, there were no significant changes in *Glut3* mRNA or protein expression/localization between groups. Western blot images show representative blots and do not include all blots/samples included in analysis. Samples were normalized across blots. Colored bars below western blots indicate sample group: teal=Control, orange= MNR, maroon= MNR+IGF1. IHC images are labeled with FC: fetal capilarries, MBS: maternal blood space, black arrows: membrane staining, scale bar: 10 μm. Control (+ sham treatment): n = 6 dams (8 female and 11 male fetuses), MNR (+ sham treatment): n = 6 dams (5 female and 11 male fetuses), MNR + IGF1: n = 5 dams (6 female and 10 male fetuses). Data are estimated marginal means ± 95% confidence interval. P values calculated using generalized estimating equations with Bonferroni post hoc analysis. *P≤0.05; **P≤0.01, ***P≤0.001

In both sexes, LAT1 staining via IHC showed an increase in intensity of nuclear/perinuclear staining in MNR placentas compared to Control, MNR+IGF1 (Indirect Exposure) and MNR+IGF1 (Direct Injection) (Figure 5A,B). *Lat2* RNA expression was unchanged between groups in males (Figure 5C, Supp. 2E). In females, *Lat2* RNA expression increased in MNR and MNR+IGF1 (Indirect Exposure) placentas compared to Control (p<0.05, p<0.05) (Figure 5D). Protein staining of LAT2 in both sexes showed a slight decrease in intensity in MNR compared to Control but an increase in staining in MNR+IGF1 (Indirect Exposure) and MNR+IGF1 (Direct Injection) compared to MNR and Control (Figure 5E,F). In males, *Snat1* expression decreased in MNR+IGF1 (Direct Injection) placentas compared to Control (p<0.05), while expression increased in MNR+IGF1 (Indirect Exposure) placentas compared to MNR and MNR+IGF1 (Direct Injection) (p<0.05, p<0.05) (Supp. 3E). *Snat1* expression did not change between groups in females (Supp. 3F). *Snat2* RNA expression decreased in MNR+IGF1 (Direct Injection) compared to Control in males (p<0.05, p<0.01) (Figure 5G, Supp. 2F) and did not change in females (Figure 5H). SNAT2 protein expression showed no changed in localization or staining intensity among groups of either sex (Figure 5I, J).

**Figure 5.**
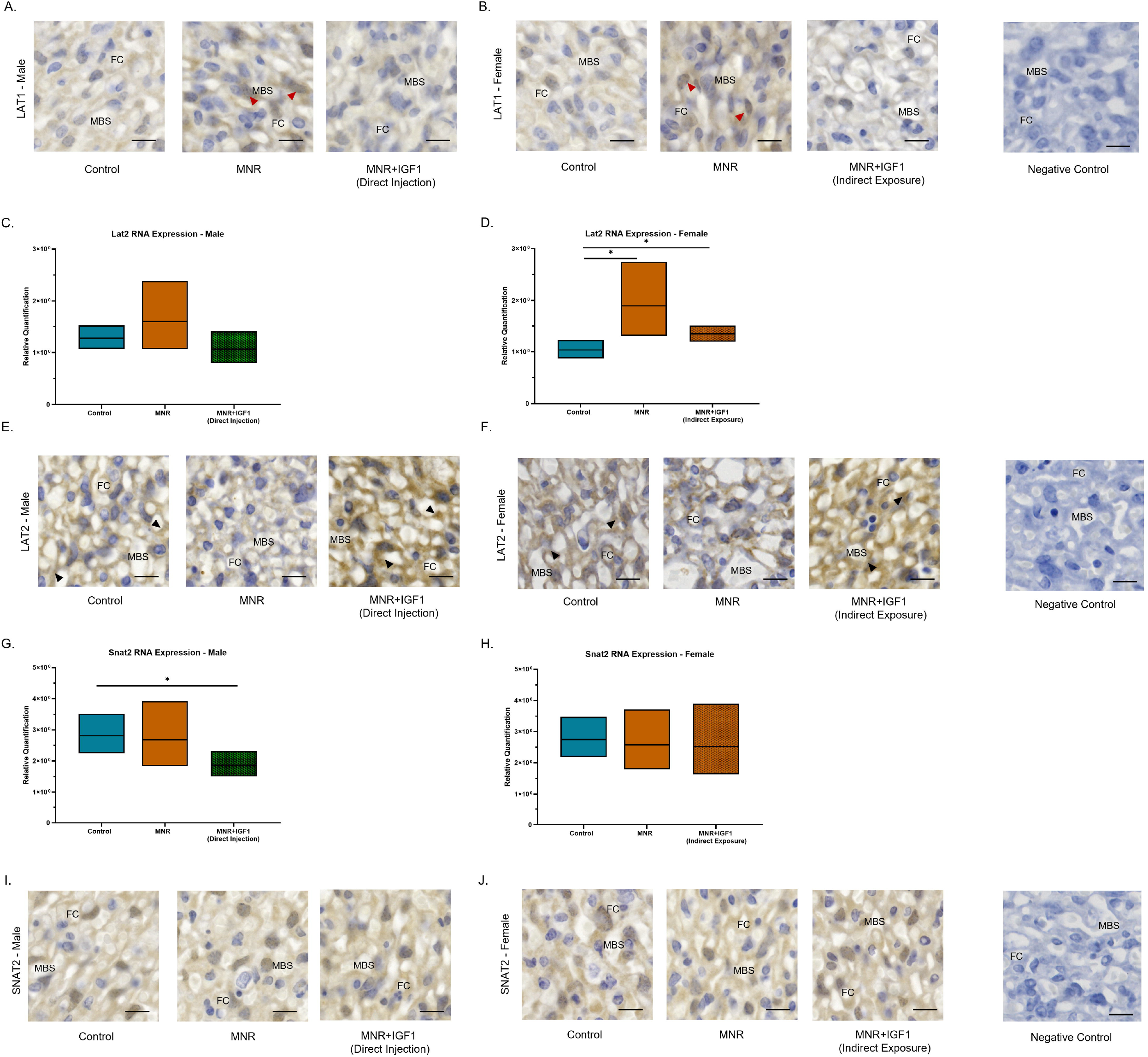
Effects of maternal nutrient restriction (MNR) and repeated *hIGF1* nanoparticle gene therapy (MNR+*IGF1*) at end of term on placental amino acid transporters. **A,B**. LAT1 IHC showed an increase in staining primarily in the nucleus in MNR compared to Control, MNR+IGF1 (Indirect Exposure), and MNR+IGF1 (Direct Injection). **C**. In males, there was no change in *Lat2* RNA expression. **D**. In females, *Lat2* RNA expression increased in MNR and MNR+IGF1 (Indirect Exposure) compared to Control. **E**,**F**. LAT2 IHC showed a decrease in staining in MNR and an increase in MNR+IGF1 (Indirect Exposure) and MNR+IGF1 (Direct Injection) compared to MNR and Control. **G**. In males, *Snat2* RNA expression decreased in MNR+IGF1 (Direct Injection) compared to control **H**. There was no change among *Snat2* placental expression in females. **I, J**. SNAT2 IHC showed no differences in staining among any groups. IHC images are labeled with FC: fetal capilarries, MBS: maternal blood space, black arrows: membrane staining, red arrows: nuclear staining, scale bar: 10 μm. Control (+ sham treatment): n = 6 dams (8 female and 11 male fetuses), MNR (+ sham treatment): n = 6 dams (5 female and 11 male fetuses), MNR + IGF1: n = 5 dams (6 female and 10 male fetuses). Data are estimated marginal means ± 95% confidence interval. P values calculated using generalized estimating equations with Bonferroni post hoc analysis. *P≤0.05; **P≤0.01, ***P≤0.001

## DISCUSSION

In the present study we aimed to elucidate the signaling changes of nutrient transport that lead to placental insufficiency/fetal growth restriction, and its mitigation with our placental nanoparticle-mediated *hIGF1* gene therapy. We demonstrated that while males and females respond to both MNR and *hIGF1* treatment similarly in overall phenotype (decreased growth in MNR and corrected growth with *hIGF1* treatment) they achieve these phenotypes through varying kinase signaling and nutrient transporter changes, as well as varying levels of *hIGF1* delivery [33]. We believe these sexual dimorphisms both shed light on the basic understanding of sex as a variable in cell/placental biology as well as hypotheses on why health outcomes vary for those affected by FGR based on fetal sex [33–35].

The importance of IGF1 in placental development and function has been well established for its role in nutrient transport [25, 26]. Our previous studies at mid pregnancy demonstrated decreased expression of *IGF1* and solute carrier genes in MNR placentas compared to Controls [32, 36]. With the addition of our placental nanoparticle-mediated *hIGF1* delivery, placental expression of positive regulators of kinase activity and phosphorylation increased [36]. Examining more of these changes at late pregnancy we now found few changes in ERK and pERK expression between either sex. AKT expression in males trended down with MNR but pAKT expression was either not altered (males) or trended upward (females) compared to controls. While there are few changes in these kinases, this is unsurprising due to the late pregnancy timepoint. Prior changes seen at mid-pregnancy were observed 5 days after placental nanoparticle-mediated *hIGF1* delivery, while in this multi-treatment study, sacrifice and tissue collection are performed 8 days after the last *hIGF1* delivery. Because of this we chose to look at more downstream effectors in the pathway from IGF1 binding to nutrient signaling.

The mTOR complex is vital for the functionality of both glucose and amino acid transporters [37, 38]. mTOR activation signals for the translocation of glucose transporters to the syncytial membrane and modulates the activity of amino acid transporters in a symbiotic relationship requiring the influx of those amino acids [38]. Interestingly, p-mTOR expression had opposing changes in the placenta dependent on sex. p-mTOR expression did not change in male MNR placentas compared to Control, while female placenta expression decreased. This opposing trend continued with *hIGF1* treatment in males showing an increase in MNR+IGF1 (Indirect Exposure), while females showed a decrease. One of the major downstream proteins from mTOR in the signaling pathway is 4EBP1, a translation initiation factor. Unsurprisingly, this protein follows the exact same changes as mTOR and p-mTOR with neither sex having differences in total protein expression, males having no change in phosphorylated 4EBP1, and females having decreased p-4EBP1 expression in MNR and MNR+IGF1 (Indirect Exposure) compared to Control. When looking at regulation of mTOR, there were few changes in positive regulators of mTOR, RICTOR and RAPTOR, while the negative regulator, DEPTOR, decreased with *hIGF1* treament from both MNR and Control in both sexes. We believe this negative regulator is downregulated in an attempt to create a homeostasis and increase p-mTOR activity, though seemingly only effective on this level in males.

Fetal growth restriction and the MNR model have been established to decrease fetal glucose transport to the fetus [8, 9, 39, 40]. We have now shown both at mid pregnancy after a single injection and at near-term after multi-treatment that our therapy is capable of increasing fetal blood glucose levels back to control, however, at near-term, females’ circulating blood glucose no longer decreased with MNR, while males were still decreased at this timepoint. This displays a mechanism in which females are able to better regulate this process, and we hypothesize this is one of the reasons female fetuses tend to fair better from FGR than their male counterparts. To elucidate why this occurred we examined RNA expression, protein expression, and protein localization of glucose transporters in the placenta. Changes were predominantly found in Glucose transporter 1 (GLUT1/SLC2a1). GLUT1 protein expression trended down in MNR compared to Control and MNR+IGF1 (Direct Injection). GLUT1 staining recapitulated this by showing more diffuse staining in MNR placentas, while Control and MNR+IGF1 (Direct Injection) placentas showed darker, defined membrane staining. Female placentas, however, showed no change in overall expression of GLUT1 among groups, while staining showed similar results as males, indicating that GLUT1 is being translated equally among females, but failing to traffic as much protein to the membrane. Other glucose transporters such as GLUT3, GLUT8, and GLUT9 had few to no changes, indicating to us that these are not the primary glucose transporters affected by FGR/MNR at this time point in the guinea pig placenta.

Amino acid transport within the placenta is primarily through system L (LAT1/2) and A (SNAT1/2) transporters [41–43]. System L-amino acid transporters are vital for many essential amino acids through sodium independent transport, while system A-amino acid transporters are essential for short chain amino acids and are sodium dependent [44, 45]. While we observed no sexual dimorphisms in protein expression among system L- or A-amino acid transporters, *Lat2* RNA expression changed in across groups in female placentas, with males having no change, and *Snat2* RNA expression changed across groups in male fetuses, with females having no change. One limitation of this study was our inability to accurately measure circulating amino acids. Because amino acids are being produced by the mother and fetus, tracer elements are needed to differentiate between the amount of amino acids transported from mother to fetus through the placenta versus the amount produced by the fetus [46–49]. In future studies we hope to perform these experiments to elucidate changes in amino acid production and transport across the placenta.

The biggest limitation of this study was our lack of directly injected female placentas. We have previously shown that all placentas in the litter express the plasmid-specific *hIGF1* following a single nanoparticle-mediated *IGF1* injection into one placenta [24], and chose to continue this approach. Fetal sex, however, cannot reliably be determined via ultrasound at mid pregnancy prior to injection time, hence we cannot control for which sex gets directly treated. While this excluded a directly injected IGF1 female group, we do show much data into females receiving lower levels of *hIGF1* via indirect exposure, which elucidate improvements to signaling, structure, and growth. Studies in non-human primates which carry a singleton pregnancy are also underway to assess translational aspects including dose, delivery, safety, and sexual dimorphisms before moving into clinical applications.

In the present study we have shown a comprehensive view of the signaling changes relevant to IGF1 and nutrient transport signaling at the end of pregnancy in the MNR/FGR placenta and with the addition of our placental nanoparticle-mediated *hIGF1* gene therapy. We show sexually dimorphic mechanisms of kinase signaling and nutrient transporter expression that we believe are critical for the understand of basic biology of sex and how it leads to our understanding of disease and potential treatment. By understanding sometimes subtle sex differences of molecular biology we hope to provide a gene therapy that is well characterized for future clinical use. This work combined with our previous studies demonstrates the potential of this therapy for future human translation and an overall healthier human population.

---

**Supplemental Figure 1.** In males, MNR+IGF1 (Indirect Exposure) had no change in protein expression of **A**. Total ERK, **B**. pERK, **C**. total AKT, **D**. pAKT, or **E**. mTOR compared to Control or MNR. **F**. p-mTOR expression increased in MNR+IGF1 (Indirect Exposure) compared to Control and MNR. **G**. RICTOR and **H**. RAPTOR expression did not change. **I**. DEPTOR expression decreased in MNR+IGF1 (Indirect Exposure) compared to Control and MNR. **J**. Total 4EBP1 had no change in protein expression. **K**. p-4EBP1 expression decreased in MNR+IGF1 (Indirect Exposure) from MNR. Control (+ sham treatment): n = 6 dams (8 female and 11 male fetuses), MNR (+ sham treatment): n = 6 dams (5 female and 11 male fetuses), MNR + IGF1: n = 5 dams (6 female and 10 male fetuses). Data are estimated marginal means ± 95% confidence interval. P values calculated using generalized estimating equations with Bonferroni post hoc analysis. *P≤0.05; **P≤0.01, ***P≤0.001

**Supplemental Figure 2.** In males, **A**. *Glut1* expression in MNR+IGF1 (Indirect Exposure) increased compared to MNR and trended upward from Control **B**. There was no change in GLUT1 expression in MNR+IGF1 (Indirect Exposure) compared to Control or MNR. **C**. *Glut3* RNA expression did not change in MNR+IGF1 (Indirect Exposure) compared to Control or MNR **D**. GLUT3 protein expression decreased in MNR+IGF1 (Indirect Exposure) compared to Control. **E**,**F**. *Lat2* and *Snat2* expression did not change among groups. Control (+ sham treatment): n = 6 dams (8 female and 11 male fetuses), MNR (+ sham treatment): n = 6 dams (5 female and 11 male fetuses), MNR + IGF1: n = 5 dams (6 female and 10 male fetuses). Data are estimated marginal means ± 95% confidence interval. P values calculated using generalized estimating equations with Bonferroni post hoc analysis. *P≤0.05; **P≤0.01, ***P≤0.001

**Supplemental Figure 3. A, B**. *Glut8* and **C, D**. *Glut9* RNA expression did not change between any groups among either sex. **E**. *Snat1* expression decreased in MNR+IGF1 (Direct Injection) compared to Control and MNR+IGF1 (Indirect Exposure), while MNR+IGF1 (Indirect Exposure) increased compared to MNR in males. F. *Snat1* expression did not change between groups in females. Control (+ sham treatment): n = 6 dams (8 female and 11 male fetuses), MNR (+ sham treatment): n = 6 dams (5 female and 11 male fetuses), MNR + IGF1: n = 5 dams (6 female and 10 male fetuses). Data are estimated marginal means ± 95% confidence interval. P values calculated using generalized estimating equations with Bonferroni post hoc analysis. *P≤0.05; **P≤0.01, ***P≤0.001

Supplemental Table 1. List of antibodies used for western blot and immunohistochemistry assays.

Supplemental Table 2. List of primers sequences used for quantitative PCR.

