## Supplementary figures and images for "Placental Nutrient Transport and Signaling in a Guinea Pig Model of Fetal Growth Restriction with Repeated Placental Nanoparticle-mediated IGF1 Treatment"

### Supplemental Figure 1

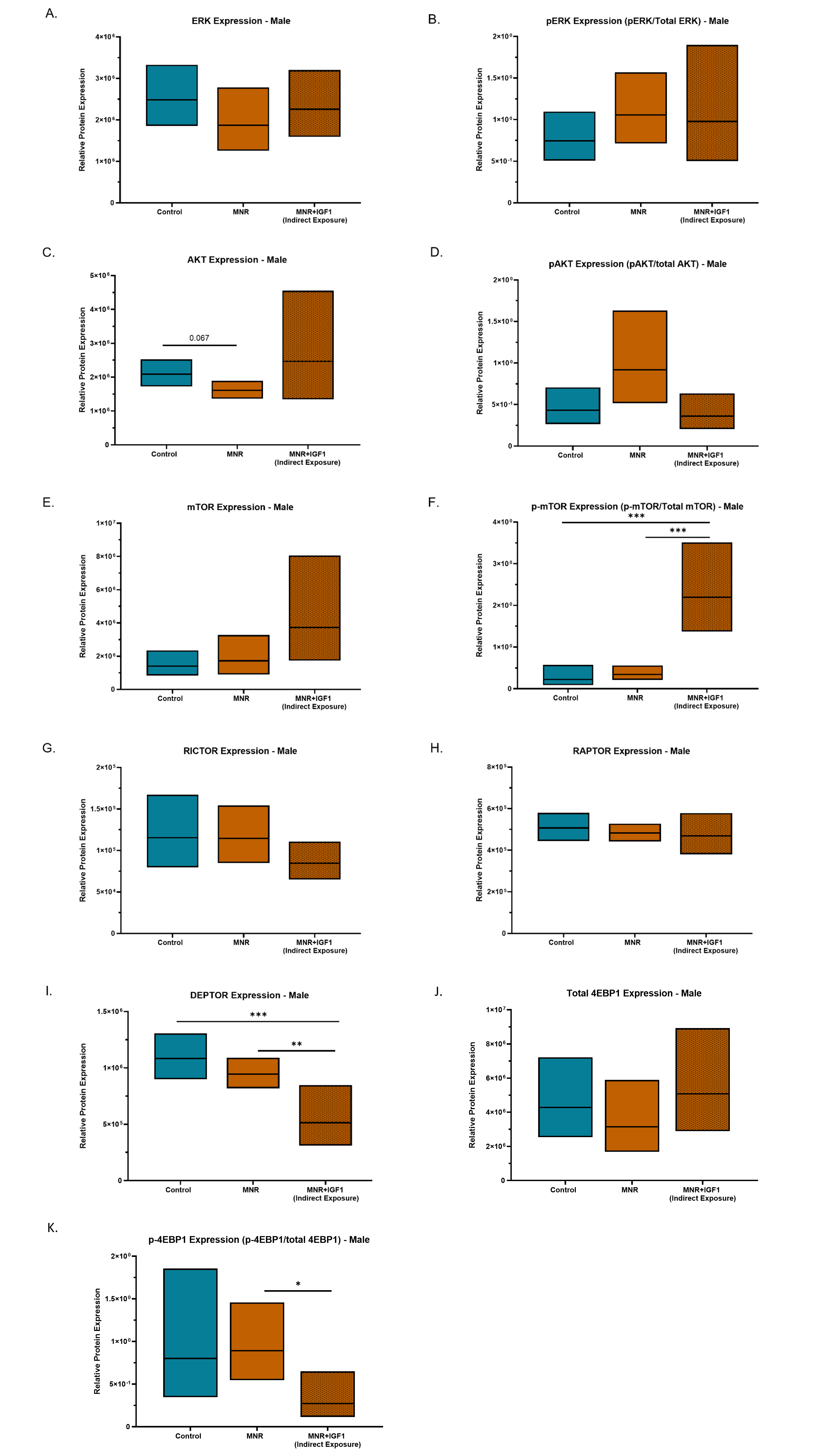

### Supplemental Figure 2

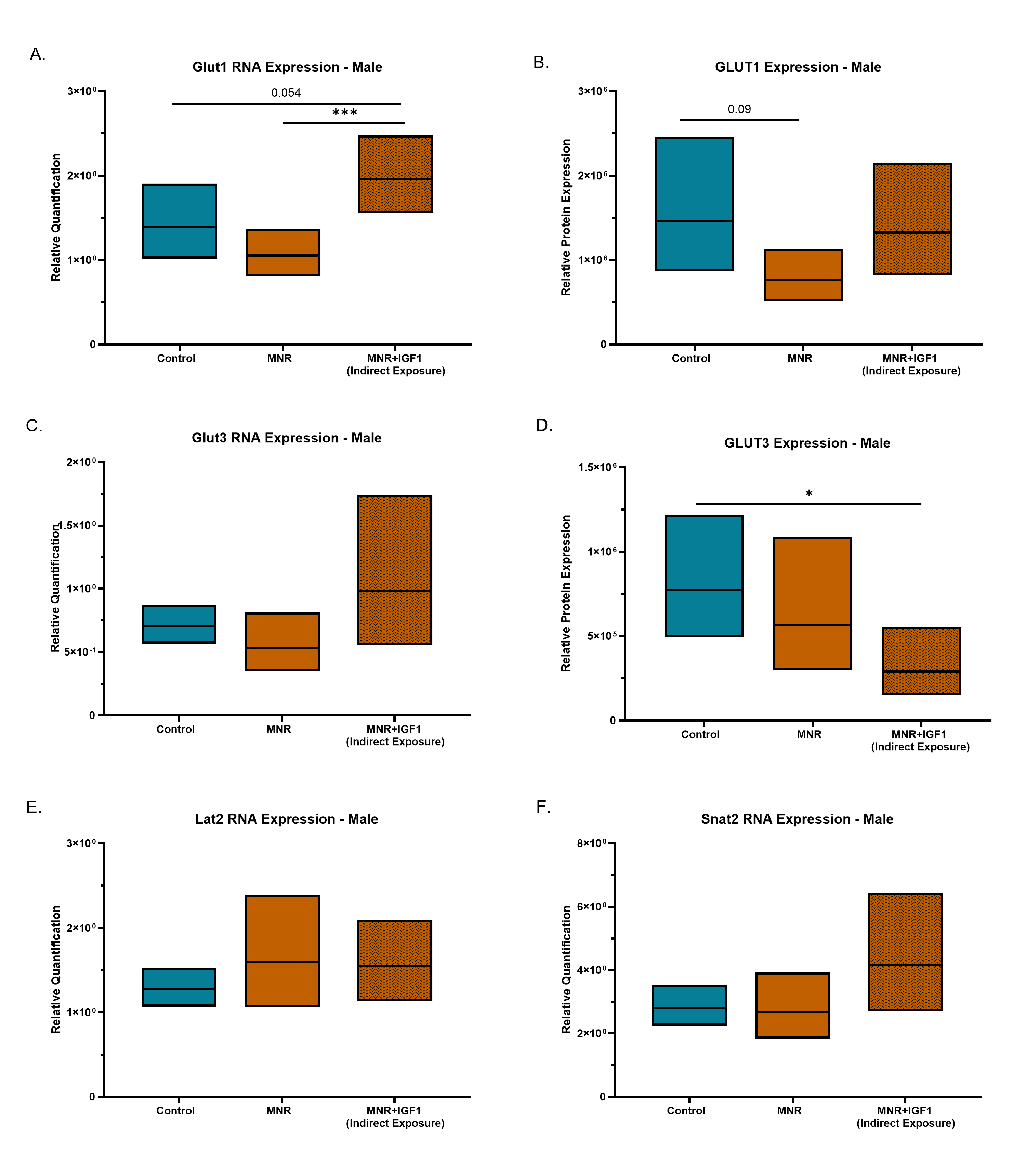

### Supplemental Figure 3

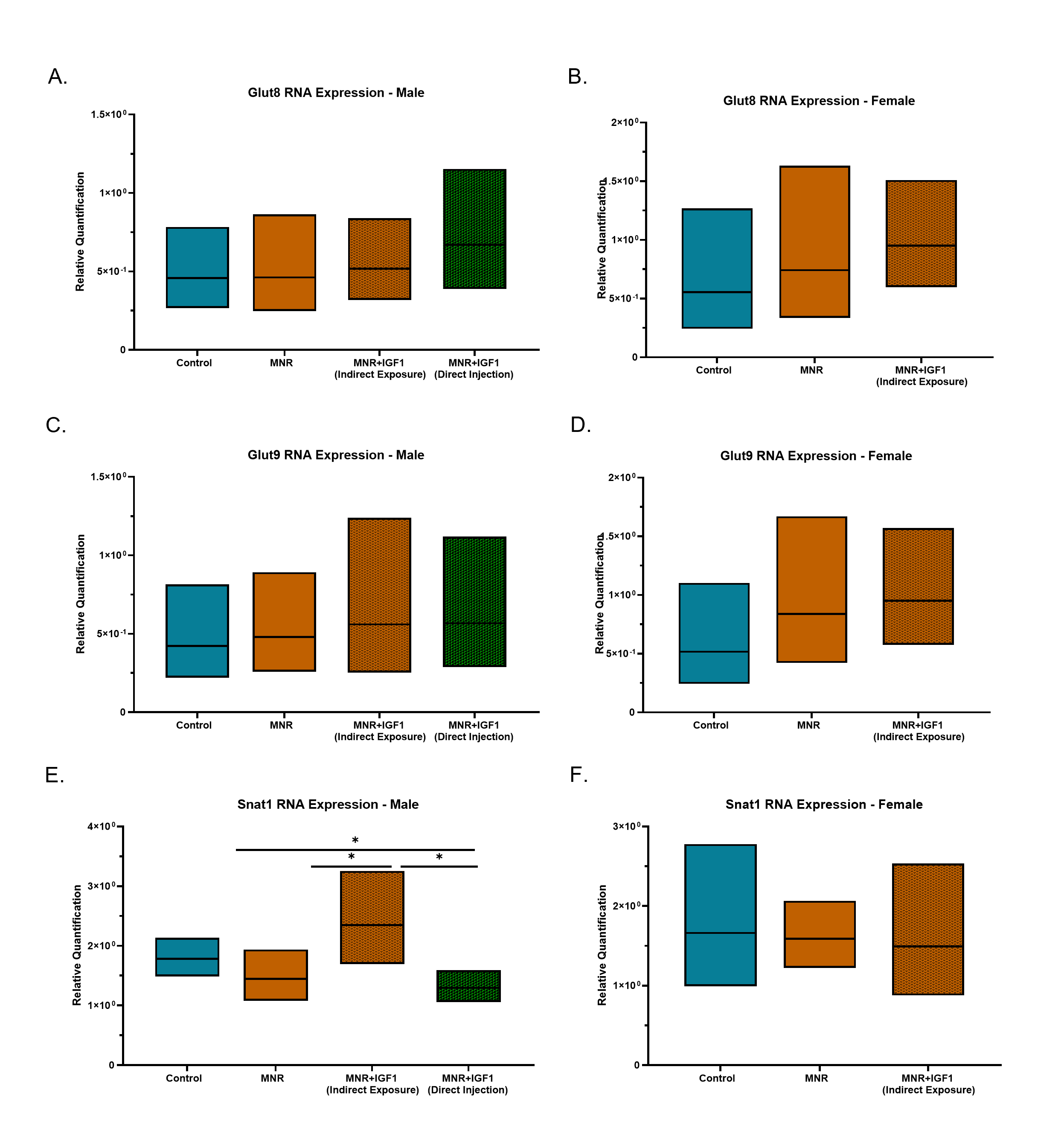

### Supplemental Table 1

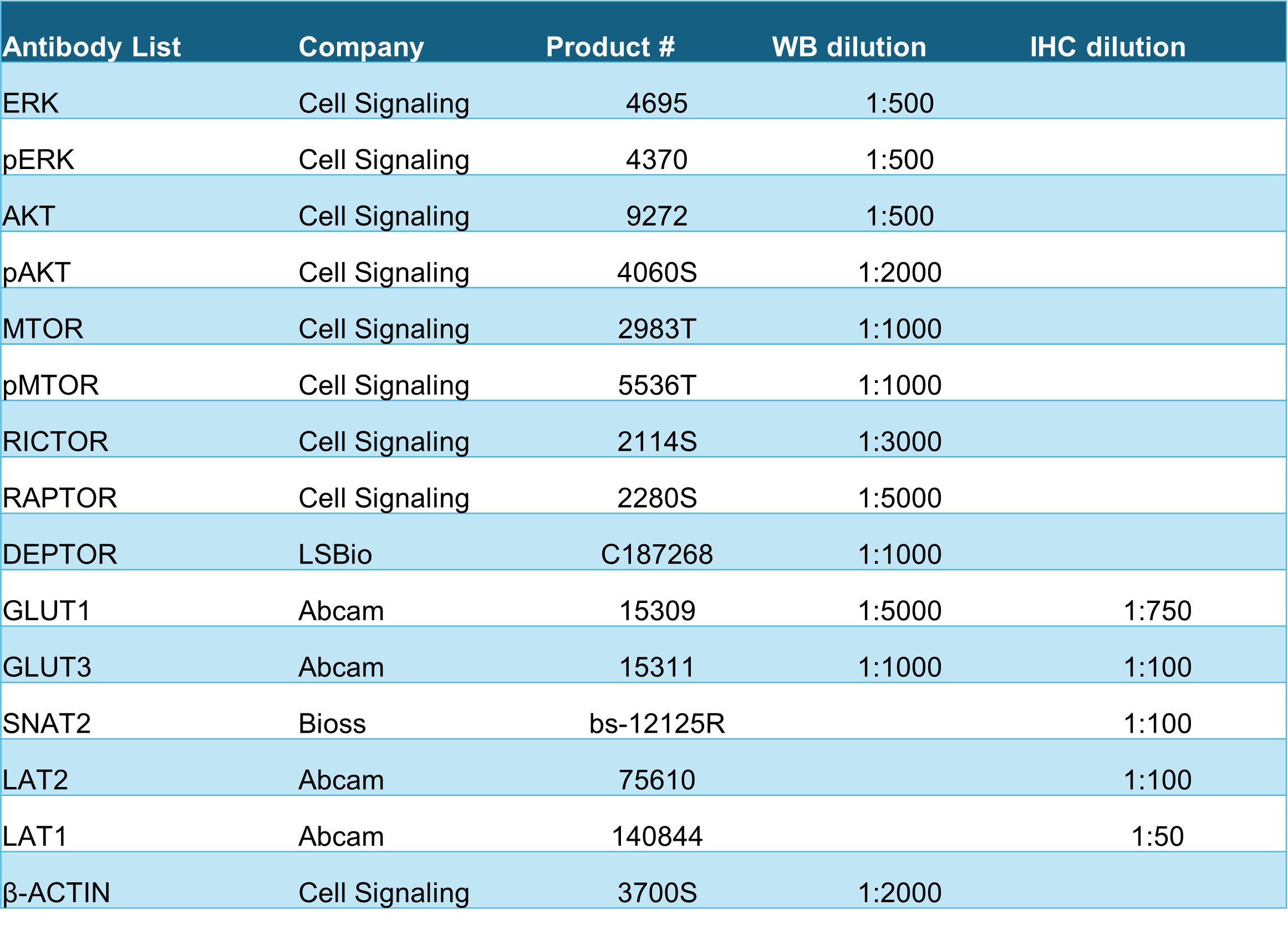

### Supplemental Table 2

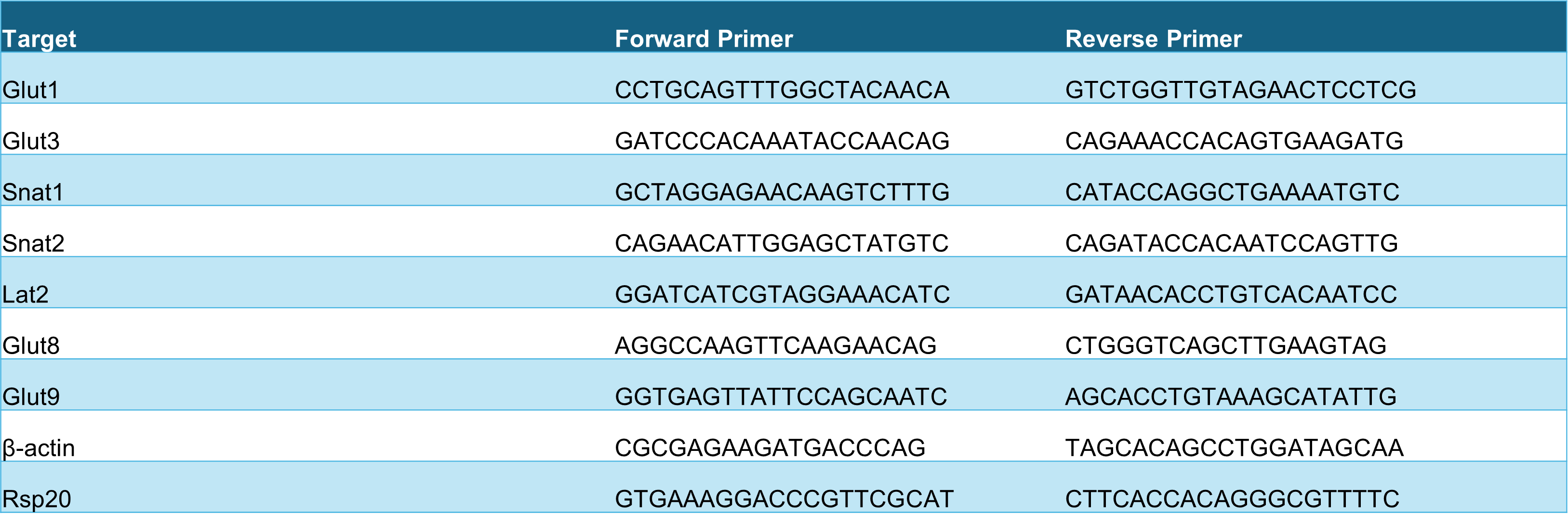
